# Variation in neotropical river otter (*Lontra longicaudis*) diet: Effects of an invasive prey species

**DOI:** 10.1101/644203

**Authors:** Diego Juarez Sanchez, John G. Blake, Eric C. Hellgren

**Affiliations:** Department of Wildlife Ecology and Conservation, University of Florida, Gainesville, Florida United States of America

## Abstract

Due to human activities, some species have expanded their distribution into areas that were historically difficult or impossible to reach by natural dispersal. Such species may become invasive if they successfully establish reproductive populations. Predation is one of the main barriers that exotic species may face in newly colonized areas. We evaluated the effect of an invasive prey (armored catfish: *Pterygoplichtys* sp.) on the dietary niche breadth and trophic level of a native predator (Neotropical river otter: *Lontra longicaudis*) in northern Guatemala. We examined otter scats from three rivers: two where the invasive armored catfish occurred and one without the invasive fish. Samples were collected two and seven years after the first report of the catfish in the area. We performed gross scat analysis and stable isotope analyses of nitrogen and carbon of fecal matter. Where the invasive armored catfish occurred, it was the main prey item for *L. longicaudis*. Particularly in the river outside of protected areas seven years after the first report of the catfish, where it accounted for 49% of the otter diet. Concordance was found between the two techniques to estimate dietary niche breadth and trophic level. The dietary niche breath of otters was narrower seven years after the invasion in comparison to two years after the invasion in both invaded rivers, but, the extent of the reduction was less inside the protected area. Finally, the trophic level of otters also showed a reduction related to the occurrence of the armored catfish on their diet.

**Resumen:** Como producto de las actividades humanas algunas especies han expandido su distribución hacia áreas que históricamente eran difícil o imposible de alcanzar mediante de dispersión natural. Estas especies pueden convertirse en invasoras si establecen exitosamente poblaciones reproductivas. La depredación es una de las principales barreras que las especies exóticas deben afrontar en las áreas recientemente colonizadas. Evaluamos los efectos de una especie invasora (el pez diablo: *Pterygoplichtys* sp.) sobre la amplitud de nicho alimenticio y el nivel trófico de un depredador nativo (la nutria de rio Neo-tropical: *Lontra longicaudis*) en el norte de Guatemala. Examinamos las excretas de nutrias provenientes de tres ríos: dos donde el pez diablo se encuentra presente y uno donde este invasor aún está ausente. Las muestras fueron colectadas dos y siete años después del primer reporte de del pez diablo en le área. Realizamos un análisis macroscópico de las excretas y análisis de isotopos estables de nitrógeno y carbono de la materia fecal. Donde el pez diablo invasor estaba presente, fue el principal ítem alimenticio de *L. longicaudis*. Particularmente en el río ubicado fuera de áreas protegidas siete años después del primer reporte del pez diablo, donde este consistió en el 49% de la dieta de la nutria. Encontramos concordancia entre las dos técnicas para estimar la amplitud de nicho dietario y nivel trófico. La amplitud de nicho dietario de las nutrias fue más angosto siete años después de la invasión en comparación con dos años luego de la invasión en ambos ríos invadidos, pero la magnitud de la reducción fue inferior dentro del área protegida. Finalmente, observamos una reducción en el nivel trófico de las nutrias relacionada con la ocurrencia del pez diablo en su dieta.

## Introduction

Predators may change their diet after an exotic prey species becomes established and abundant in the predator’s range [1–4]. Inclusion of such a species in a predator’s diet can lead to a shift in the predator’s dietary niche, which may become wider or narrower, depending on the intensity of use of the new resource and changes in the use of alternative native prey [5]. Furthermore, the type of prey that a predator eats defines its trophic level (e.g., primary consumer, secondary consumer). Both niche breadth and trophic levels can be evaluated using gross scat analysis and stable isotopes analyses.

An important group of invasive species in freshwater communities are the armored catfishes of the South American family Loricariidae, a diverse group of fishes with 928 valid species and eight subfamilies, including the genus *Pterygoplichthys*, commonly known as the suckermouth armored catfish (hereafter ACF; [6]). These catfish are very popular in the aquarium trade, easily domesticated, exhibit parental care [7], possess physiological tolerance to adverse conditions [8–12], have wide distribution ranges [13], and possess high reproductive and growth rates [14,15]. They feed on detritus, an abundant resource, especially in human-modified areas, and therefore have a low fractional trophic level (FTL) [13]. These traits contribute to their invasiveness, as they fulfill the six life-history variables associated with species that successfully establish invasive populations [16]. The presence of ACF as an invasive species has been documented for at least 21 countries in five continents [17]. In 2005, an established population of *Pterygoplichthys pardalis* was found in Laguna Frontera at the mouth of the Usumacinta River, Tabasco, Mexico [18]. Two years later, *P. pardalis* was reported in Guatemala in the headwaters of the San Pedro River, a tributary of the Usumacinta River (Juarez-Sanchez and J. F. Moreira, in prep.). The species identification, however, has not been confirmed because *P. pardalis* can be misidentified and confused with other species of *Pterygoplichthys* given that identification is based on ventral color patterns and hybridization with *P. disjunctivus* has been reported elsewhere [19–21].

The ACF has been reported to have positive effects by generating nutrient hotspots, making nutrients available for producers in nutrient-depleted areas [22]. However, the amount of nutrients released by the ACF does not compensate for its grazing pressure [23]. Other negative impacts of ACF have been documented in places where they have established invasive populations. These impacts include asphyxiating native predators in Puerto Rico [24]; preying on native fish eggs and first-feeding fry in Thailand [25]; competing for forage with native species, reducing biofilm from the substrate, and changing the proportions of dissolved nutrients in the Philippines and Mexico [23,26,27]; harassing manatees [28–30]; and possibly promoting erosion with their nesting burrows in Florida and Mexico [7,31]. These impacts could occur anywhere ACF establish an invasive population. Invasive ACF are preyed upon by native piscivorous predators such as common snook (*Centropomus undecimalis*) and the Neotropical cormorant (*Phalacrocorax brasilianus*) [32,33], although their effects on these and other native predators have not been evaluated.

Otters (Lutrinae) are mid-sized carnivores that are top predators in freshwater wetlands and riverine systems because of their high energetic demand and trophic position [34,35]. The Neotropical river otter (*Lontra longicaudis*; hereafter NRO) is a semi-aquatic mustelid that preys primarily on benthic slow-moving fish [36], but also feeds on crustaceans, mollusks, reptiles, and mammals (Table 1). This species is distributed from northern Mexico to northern Argentina, coexisting with different community assemblages of prey species, and adapting its foraging behavior according to the local community [37]. Where ACF are native, they coexist with the NRO and constitute one of the most important prey items in its diet [37–40]. However, the role of ACF as a prey item for NRO in areas where ACF has been introduced is unknown and may be reshaping the foraging ecology of the NRO in those areas.

**Table 1.** Food items reported as present in diets of Neotropical river otters across their geographic range.

| Locality | Primary item | Other items | Citation |
| --- | --- | --- | --- |
| Oaxaca, México. | crustaceans (53.0%) | fish (33.1%), insects (9.8%) and amphibians (4.0%) | [41] |
| México state, México. | fish (92.4%) | invertebrates (3.5%), amphibians (2.9%) and plant matter (1.8%) | [42] |
| Alto Cauca, Colombia. | fish (76.7%) | insects (12.7%), reptiles (0.7%), and others (9.9%) | [39] |
| Salta, Argentina. | fish (53%) | insects (24%), crustaceans (16%), amphibians (7%), and reptiles, mammals and mollusks (<0.1%) | [37] |
| Rio de Janeiro, Brazil. | fish (86%) | crustaceans (71%), amphibians (10%), mammals (3%), birds (0.6%), reptiles (0.2%) and others (0.7%) | [43] |
| Rio Grande do Sul, Brazil. | Fish (Loricariidae 41.1%, Cichlidae 21%, Pimelodidae 12.6%, Characidae 6.5 %) | other fish (12.5 %), Megaloptera (3.6%), mammals (1.2%), insects (0.4%), Decapoda (0.1%), birds (0.3%), snakes (0.3%) and plant matter (0.4%) | [38] |
| Rio Grande do Sul, Brazil. | fish (82.6 %) | crustaceans (20.6%), birds (4.5%), mammals and snakes (3.7%), Coleoptera (1.2%), amphibians (0.8%) and mollusks (0.4%) | [44] |
| Rio Grande do Sul, Brazil | fish | mammals, amphibians, birds, snakes, insects, crustaceans mollusks and eggs. | [45] |

Percent values are frequency of occurrence and do not add to 100%.

The main objective of this study was to determine if invasive armored catfish affected the diet of Neotropical river otters. Given that NRO feed on ACF in areas where native populations overlap [38,46,47], we hypothesized that NRO will change their diet to include ACF in rivers where invasive populations of ACF occur. We predicted that where ACF are present, they will become the main prey of NRO and reduce the niche breadth of NRO. If ACF become the main prey of the NRO, we also predicted a lower trophic level for the NRO in areas where ACF are present due to the low trophic level of the ACF.

## Materials and methods

### Study area

The study area is located at northern Guatemala in the district of Petén (between 15.50° and 17.50° N and −88.50° and −91.25° W) and includes the Usumacinta and Mopan rivers (Fig. 1). Precipitation ranges from 1,200 to 4,000 mm/year on a gradient decreasing northward (INSIVUMEH, 2016). Major habitat types in the study area consist of subtropical moist forest in the north, subtropical very moist forest in the south, and tropical very moist forest in the southeast [48]. The entire study area consists of lowland forest, with elevations ranging from 0 to 1000 masl.

**Fig 1.**
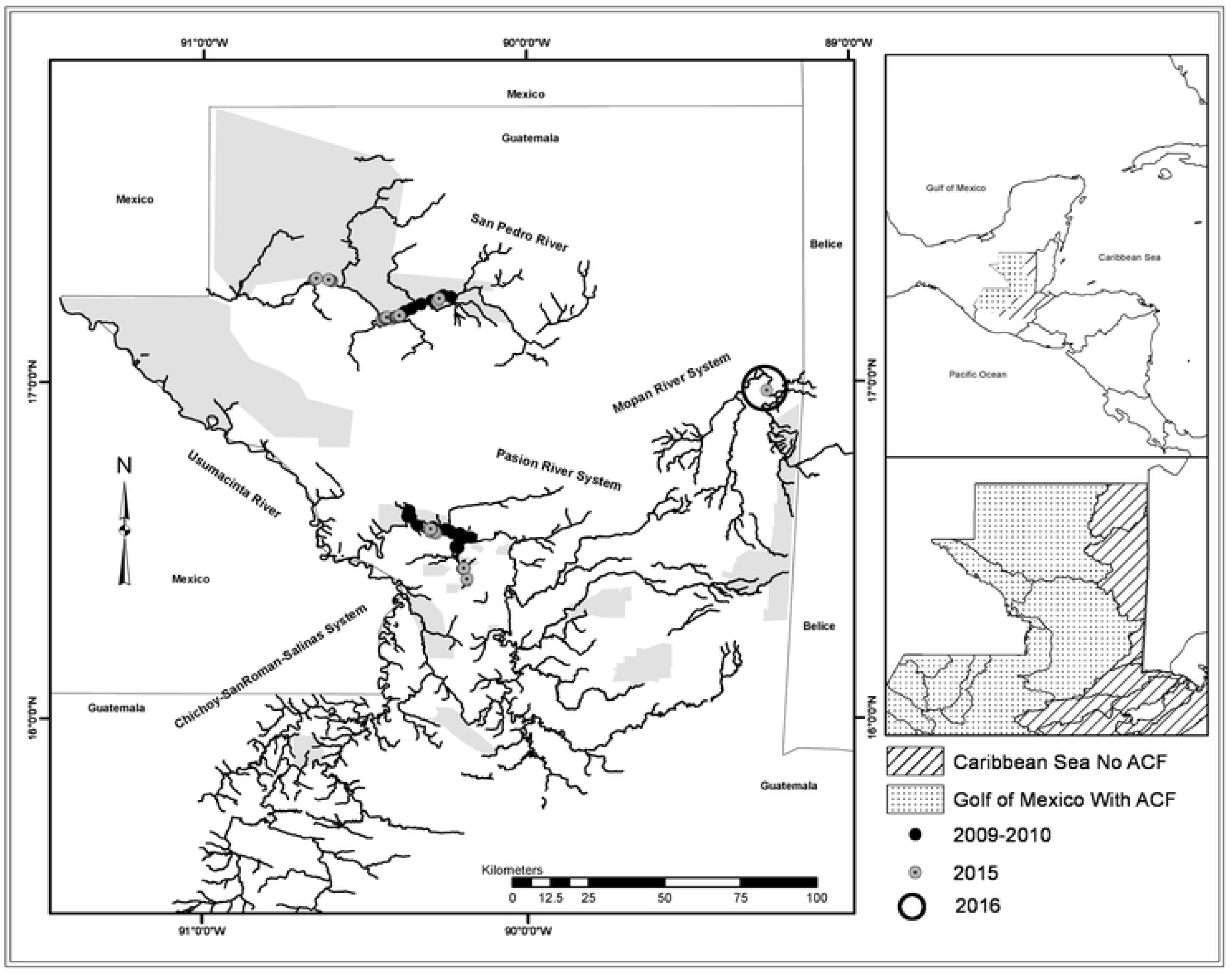
**Study area** for collection of Neotropical river otter scats in northern Guatemala. Grey circles represent samples collected in 2009-2010; black solid circles represent samples collected in 2015; black hollow circle represents the area where samples were collected in 2016. The dashed area represents the Usumacinta basin divided in sub-basins, where the armored catfish has been reported (ACF). The striped area represents the Caribbean runoff where no ACF has been reported. Grey areas represent protected areas.

In northern Guatemala, rivers flow into the Gulf of Mexico or into the Caribbean Sea watersheds (Fig. 1). Thus, bodies of fresh water are isolated by large expanses of land in the headwaters, and large distances between river mouths along the coast. The Mopan River flows northwards from southern Petén and then east in central Belize into the Caribbean Sea. The Usumacinta River runs northwest into the Gulf of Mexico. Samples were collected from the Mopan River and two tributaries of the Usumacinta River: The San Pedro River and the Pasion River. In Guatemala, the San Pedro River flows along the southern border of Laguna del Tigre National Park whereas the Pasion and the Mopan rivers mainly run through private lands that are under different land uses.

The Usumacinta basin has at least 61 fish species distributed in 25 families. The two main families in Usumacinta basin are Cichlidae with 18 species and Poeciliidae with 10 species [6,13,49,50]. To our knowledge, no peer-reviewed document has been published that describes fishes of the Mopan River within the borders of Guatemala. Thus, information about the fish assemblage in this river is based on information from the estuarine area in Belize. Therefore, the number of fish species that we are considering as present and potential prey for otters in the river headwaters within Guatemalan territory may be inflated. In Mopan River, there are at least 103 fish species distributed in 32 families, including the invasive tilapia (*Oreochromis aureus*). The main families are Cichlidae with 14 species and Poeciliidae with 16 species [6,13,51]. Exotic tilapia is widespread across all Guatemala due to multiple introductions, both accidental and deliberate from aquaculture or governmental fisheries restocking. The Asian grass carp (*Ctenopharyngodon idella*) and the ACF have been found in the Usumacinta basin, but the origins of these invasions are not clear.

### Scat Collection

Samples were collected during three periods: May 2009 – April 2010, May – July 2015, and June 2016. The search for otter scats was conducted from a small boat moving at slow speeds (< 5 km/h) close to the shoreline, with scats and latrines typically found on protruding structures (e.g. rocks or fallen trees). This search was conducted along both shorelines of the river in opposite directions. All scats were collected, placed in paper and/or plastic bags with silica gel, and stored in a dry environment. Otter scats were identified by their appearance, as no other species present in the study area have similar scats (located on protruding sites along the river shore, low fecal matter and high content of fish or crab remains) [52]. If a scat was found but its identification was doubtful, it was collected and included in the analysis only if otter hair from grooming was found on it. Each scat was assigned a unique code and the geographic coordinates of its location were recorded using a handheld GPS unit (GARMIN © Astro 320, Garmin Ltd. Kansas City, USA).

We sampled the Usumacinta basin during 2009-2010 using continuous searches along the rivers, including 38.5 km of the San Pedro River (starting from Paso Caballos village and moving west) and 89.1 km of the Pasion River (starting from Sayaxche town and heading west). We sampled the Usumacinta and Mopan basins in 2015 by organizing the search for scats into segments of 10 km, with segments separated by at least 10 km. In the Usumacinta basin, we sampled along 40, 50, and 30 km of the San Pedro, La Pasion, and Usumacinta rivers, respectively. Surveys began in Paso Caballos for the San Pedro, in Sayaxche for the Pasion, and in Betel town for the Usumacinta River. The Mopan River was sampled along 10 km in 2015 near La Polvora military base. In June 2016, local fisherman collected samples for in the Mopan River near La Polvora military base, no exact georeference was collected per sample (Fig. 1).

### Scat Sample Preparation and Analysis

Samples of fecal matter were collected from each scat, homogenized using a porcelain mortar and pestle, stored in glass vials and sent to the Light Stable Isotope Mass Spectrometry Laboratory in the Department of Geological Sciences at the University of Florida for stable isotope analysis (SIA) of δ^15^N and δ^13^C. Samples were analyzed using a Thermo Electron DeltaV Advantage isotope ratio mass spectrometer coupled with a ConFlo II interface linked to a Carlo Erba NA 1500 CNHS Elemental Analyzer. All carbon isotopic results are expressed in standard delta notation relative to VPDB. All nitrogen isotopic results are expressed in standard delta notation relative to AIR. Hard remains (i.e., scales, skeleton pieces) were separated and identified to the lowest possible taxonomic level. A list of potential prey species for otters was made, consisting of all the fish species reported in the study area that have a reported maximum total length ≥ 100 mm (S1 Table). Size selection was based on the assumption that otters prefer to feed on fish within the 100-150 mm size range [53]. Prey remains that could be identified were fish scales, otoliths or vertebra; crustacean shells; and mammal hairs. A scale guide was constructed for 68 of the 80 scaled fish species that are found in the sampled river basins and that were consider potential prey of the NRO [54]. Scales were obtained from museum specimens at the Florida Museum of Natural History (FLMNH) and El Colegio de la Frontera Sur in México (ECOSUR). Scales from these fish species were cleaned with water and alcohol, placed on glass slides with nail polish, and sealed with a coverslip to make semi-permanent slides. For 10 catfish species that do not have scales, the identification was based on fin spines, using reference material from the zooarchaeological collection at FLMNH. Hairs found in the scats were identified using a hair-identification guide [55] and reference material from the mammal collection of the Museo de Historia Natural (MUSNAT) at the Universidad de San Carlos de Guatemala (USAC). Otter hair (product of grooming) was saved and pressed between glass slides and coverslips for future analysis.

### Data Analyses

For data analyses, the sampling units were the rivers (San Pedro, La Pasion, Mopan) with year as factor (2009-2010 and 2015-2016). The year effect represents 2 and 7 years after the advent of the ACF invasion. Comparisons were made over time (i.e., same river, different year) only using data from Pasion and San Pedro rivers where the ACF are present; we additionally looked at differences across river basins in the same sampling years (i.e., different river, same year), combining 2015-2016 records as one year and including the Mopan River where ACF do not occur.

The importance of each prey species can be biased by abundant and conspicuous hard remains that are identifiable for some species, even if those species are consumed in low numbers, due to differential digestibility of prey items. This overestimation of some species can then lead to an underestimate of overall diet diversity. On the other hand, when using SIA of predators, one can measure diet diversity breadth and comparative trophic levels but with no taxonomic information about the prey. For this reason, we used both techniques, expecting to find concordance between them.

#### Gross scat analysis (GSA)

Accumulation curves were constructed using program EstimateS (© Colwell 2013, Connecticut, USA) where the expected number of prey species found in a given number of scats is obtained by

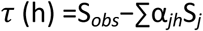

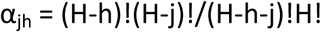

were τ (h) is the estimated number of species for h number of scats; S_*obs*_ is the number of species actually observed; S_*j*_ is the number of prey species found in j scats; α_*jh*_ is a combinatorial coefficient; H is the total number of scats; h is the number of possible combination of scats that add up to j scats; and j is the number of scats per moment or segment of the curve [56].

The importance of different food items, including the ACF, in the NRO diet was assessed through GSA, using the percentage of occurrence. Percentage of occurrence was estimated for a prey item by dividing the number of scats with item *i* by the total number of reported items. To compare the NRO niche breadth between basins, with different prey communities, Levin’s index was used:

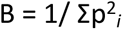

where p is the proportion of food items from category *i* [57]. The Levin’s niche-breadth index can be standardized using:

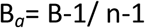

where Ba is the standardized Levin’s niche-breadth index, B is Levin’s niche-breadth index, and n is the number of recorded species. Levin’s index ranges from 1 to n and from 0 to 1 in its standardized version. In both cases, its minimum value is reached when all reported prey belongs to only one species (specialist predator) and is at its maximum when all the species are consumed in the same proportion (generalist predator). It has been suggested that values of B_*a*_ > 0.6 represent a generalist and values of B_*a*_ < 0.4 a specialist [58,59]. To estimate confident intervals the samples (scats) were randomly selected with replacement (bootstrap), then the index was re-estimated with the resulting set of samples. This procedure was repeated 1000 times and the confidence intervals calculated.

The NRO’s fractional trophic level, which represents the trophic distance of a consumer species from producers, in each basin was estimated using Pauly and Palomares’s (2005) formula:

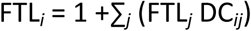

where FTL_*i*_ is the fractional trophic level of the consumer, +1 is a constant increment for the FTL of a consumer, FTL_*j*_ is the fractional trophic level of the prey *j*, and DC_*ij*_ is the proportion of contribution of prey *j* to the diet of consumer *i*. Prey FTL*j* values were obtained from FishBase database[13]. The DC_*ij*_ was based on proportion of occurrence values by river-year combination in the otter scats. To estimate confidence intervals a bootstrap procedure was developed as explained above.

#### Stable isotope analysis (SIA)

Stable isotope analysis (SIA) measures the proportion of heavy to light stable isotopes in a sample [63,64]; its values are expressed in delta notation (δ) or per mil (‰) and estimated with this equation:

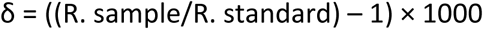

where R = heavy isotope / light isotope obtained with a mass spectrometer.

Isotopic values of a predator are higher than those of its prey due to a process called fractionation, wherein the molecules with the lighter isotopes, given their lighter overall weight, react faster and can be metabolized and excreted faster than the heavier ones. This process results in the predator being enriched with a higher proportion of heavier isotopes than its prey [61,64]. The mean value of this fractionation across taxa is 3.4‰ (1 SD = 1‰) for δ^15^N and 0.4‰ (1 SD = 1.3‰) for δ^13^C [61]. These values are the expected increment of the isotopic value when molecules are assimilated from prey tissue to predator tissue (from lower to higher trophic levels;[65]).

Isotopic values of different tissues, such as bone, blood, hair or muscle, have been used to evaluate the diets of a wide range of species [66–73]. Normally, tissue samples are obtained from dead or captured specimens but these invasive techniques are sometimes difficult or impossible to use, especially for secretive, rare or endangered species. However, controlled experiments have shown that SIA based on feces is sensitive to changes in the diet over periods of 3 hours for insectivorous bats [74] and, thus, represent the isotopic values of the latest meals of the individual that produced the scat[75]. In carnivores and omnivores, SIA based on scats can be used to estimate the main type of prey and nutrient flow, using δ^15^N to infer the range of trophic positions or FTLs at which a predator eats, and δ^13^C to determine the type of producers that supported the specific trophic chain [61,76–78]. Further, the variance of isotopic values of a population may represent the niche width (or breadth) of a consumer [79].

Taking δ^15^N and δ^13^C values from individual scats as samples from each river, we evaluated the data for normality using histograms, qq-plots and a Shapiro-Wilk normality test; all values followed a normal distribution. To evaluate differences between variances in δ^13^C and δ^15^N as a niche breadth metric, a Levene’s homoscedasticity test was used. To test for differences in mean δ^15^N values between rivers and years we use a two-factor ANOVA after a log transformation of the data to correct for lack of homoscedasticity; a post-hoc paired *t*-test with Bonferroni adjusted *p*-values was used to evaluate where the differences occurred. All the statistical procedures except for the species accumulation curves were performed using the program R [80].

## Results

Field collection of scats yielded 286 samples identified as coming from the NRO. After eliminating scats that had some type of contamination (e.g., wood, mud, or termite nest), 177 samples of fecal matter were sent for isotopic analysis (Table 2). We identified 35 scaled fish species, including three nonnative fish species (*Oreochromis aureus, Ctenopharyngodon idella* and *Pterygoplichtys* sp.) from otter scats. In addition, remains of unidentified insects, one reptile, one unidentified mammal, and unidentified crabs and crayfish were recovered from the scats (Table 3).

**Table 2.** Scats of Neotropical river otters collected in northern Guatemala.

| River | No. of scats collected<br>(year) | No. of scats without<br>contamination (year) |
| --- | --- | --- |
| Usumacinta | 1 (2015) | 0 |
|  | 1 Total | 0 Total |
| San Pedro | 36 (2009) | 20 (2009) |
|  | 117 (2015) | 55 (2015) |
|  | 153 Total | 75 Total |
| La Pasion | 52 (2010) | 36 (2010) |
|  | 40 (2015) | 34 (2015) |
|  | 92 Total | 70 Total |
| Mopan | 1 (2015) | 1 (2015) |
|  | 39 (2016) | 31 (2016) |
|  | 40 Total | 32 Total |

Only scats without contamination were used for fecal matter isotope analyses

**Table 3.** Number of records (No), and percentage of the total of prey species (%) found in otter scats collected from the Mopan, Pasion and San Pedro (San Pe.) rivers, northern Guatemala, during 2009-2010 (09-10) and 2015-2016 (15-16).

|  | Mopan<br>15-16 |  | Pasion<br>09-10 |  | Pasion<br>15-16 |  | San Pe.<br>09-10 |  | San Pe.<br>15-16 |  |
| --- | --- | --- | --- | --- | --- | --- | --- | --- | --- | --- |
|  | No | % | No | % | No | % | No | % | No | % |
| Belonidae |  |  |  |  |  |  |  |  |  |  |
| *Strongylura hubbsi* | 0 | 0 | 0 | 0 | 0 | 0 | 3 | 2.2 | 0 | 0 |
| *Strongylura marina* | 1 | 1.1 | 0 | 0 | 0 | 0 | 0 | 0 | 0 | 0 |
| Carangidae |  |  |  |  |  |  |  |  |  |  |
| *Caranx latus* | 1 | 1.1 | 0 | 0 | 0 | 0 | 0 | 0 | 0 | 0 |
| Centropomidae |  |  |  |  |  |  |  |  |  |  |
| *Centropomus ensiferus* | 1 | 1.1 | 0 | 0 | 0 | 0 | 0 | 0 | 0 | 0 |
| Characidae |  |  |  |  |  |  |  |  |  |  |
| *Astianax fasciatus* | 0 | 0 | 0 | 0 | 0 | 0 | 9 | 6.5 | 1 | 0.3 |
| Cichlidae |  |  |  |  |  |  |  |  |  |  |
| *Chuco intermedius* | 6 | 6.6 | 0 | 0 | 0 | 0 | 0 | 0 | 1 | 0.3 |
| *Cincolichthys bocourti* | 1 | 1.1 | 0 | 0 | 1 | 1.4 | 0 | 0 | 1 | 0.3 |
| *Cincolichthys pearsei* | 0 | 0 | 0 | 0 | 0 | 0 | 0 | 0 | 1 | 0.3 |
| *Cribroheros robertsoni* | 1 | 1.1 | 13 | 9.1 | 4 | 5.5 | 22 | 15.8 | 12 | 4.1 |
| *Kihmchithys ufermammi* | 0 | 0 | 0 | 0 | 0 | 0 | 3 | 2.2 | 1 | 0.3 |
| *Maskaheros argenteus* | 0 | 0 | 1 | 0.7 | 0 | 0 | 0 | 0 | 0 | 0 |
| *Mayaheros urophthalmus* | 2 | 2.2 | 5 | 3.5 | 5 | 6.8 | 5 | 3.6 | 17 | 5.9 |
| *Oreochromis aureus* | 8 | 8.8 | 5 | 3.5 | 6 | 8.2 | 4 | 2.9 | 21 | 7.3 |
| *Parachromis friedrichsthalii* | 2 | 2.2 | 9 | 6.3 | 5 | 6.8 | 1 | 0.7 | 4 | 1.4 |
| *Petenia splendida* | 0 | 0 | 1 | 0.7 | 0 | 0 | 0 | 0 | 3 | 1 |
| *Rheoheros lentiginosus* | 0 | 0 | 1 | 0.7 | 1 | 1.4 | 0 | 0 | 0 | 0 |
| *Rocio octofasciata* | 0 | 0 | 2 | 1.4 | 1 | 1.4 | 0 | 0 | 3 | 1 |
| *Thorichthys affinis* | 0 | 0 | 2 | 1.4 | 0 | 0 | 0 | 0 | 1 | 0.3 |
| *Thorichthys aureus* | 6 | 6.6 | 0 | 0 | 0 | 0 | 0 | 0 | 0 | 0 |
| *Thorichthys meeki* | 5 | 5.5 | 11 | 7.7 | 2 | 2.7 | 10 | 7.2 | 25 | 8.6 |
| *Thorichthys pasionis* | 0 | 0 | 10 | 7.0 | 1 | 1.4 | 10 | 7.2 | 8 | 2.8 |
| *Trichromis salvini* | 0 | 0 | 0 |  | 0 | 0 | 0 | 0 | 2 | 0.7 |
| *Vieja bifasciata* | 0 | 0 | 3 | 2.1 | 6 | 8.2 | 1 | 0.7 | 21 | 7.3 |
| *Vieja melanurus* | 0 | 0 | 3 | 2.1 | 1 | 1.4 | 0 | 0 | 8 | 2.8 |
| Cyprinidae |  |  |  |  |  |  |  |  |  |  |
| *Ctenopharyngodon idella* | 0 | 0 | 7 | 4.9 | 0 | 0 | 0 | 0 | 5 | 1.7 |
| Eleotridae |  |  |  |  |  |  |  |  |  |  |
| *Dormitator maculatus* | 1 | 1.1 | 0 | 0 | 0 | 0 | 0 | 0 | 0 | 0 |
| Gerreidae |  |  |  |  |  |  |  |  |  |  |
| *Eugerres mexicanus* | 0 | 0 | 0 | 0 | 0 | 0 | 0 | 0 | 4 | 1.4 |
| Hemiramphidae |  |  |  |  |  |  |  |  |  |  |
| *Hyporhamphus mexicanus* | 0 | 0 | 0 | 0 | 0 | 0 | 9 | 6.5 | 2 | 0.7 |
| Lepisosteidae |  |  |  |  |  |  |  |  |  |  |
| *Aractosteus tropicus* | 0 | 0 | 0 | 0 | 0 | 0 | 1 | 0.7 | 0 | 0 |
| Loricariidae |  |  |  |  |  |  |  |  |  |  |
| *Pterygoplichthys* sp | 0 | 0 | 14 | 9.9 | 36 | 49.3 | 23 | 16.5 | 75 | 25.9 |
| Megalopidae |  |  |  |  |  |  |  |  |  |  |
| *Megalops atlanticus* | 0 | 0 | 0 | 0 | 0 | 0 | 0 | 0 | 1 | 0.3 |
| Mugilidae |  |  |  |  |  |  |  |  |  |  |
| *Mugil cephalus* | 0 | 0 | 1 | 0.7 | 0 | 0 | 0 | 0 | 0 | 0 |
| Poeciliidae |  |  |  |  |  |  |  |  |  |  |
| *Belonesox belizanus* | 0 | 0 | 5 | 3.5 | 0 | 0 | 1 | 0.7 | 12 | 4.1 |
| *Poecilia mexicana* | 1 | 1.1 | 3 | 2.1 | 1 | 1.4 | 17 | 12.2 | 24 | 8.3 |
| *Poecilia petenensis* | 0 | 0 | 11 | 7.7 | 1 | 1.4 | 17 | 12.2 | 26 | 9 |
| Ariidae, Heptapteridae, Ictaluridae |  |  |  |  |  |  |  |  |  |  |
| Catfish | 8 | 8.8 | 3 | 2.1 | 1 | 1.4 | 3 | 2.2 | 4 | 1.4 |
| Crab | 32 | 35.2 | 30 | 21.1 | 0 | 0 | 0 | 0 | 3 | 1 |
| Crayfish | 0 | 0 | 0 | 0 | 0 | 0 | 0 | 0 | 1 | 0.3 |
| Insect | 3 | 3.3 | 0 | 0 | 0 | 0 | 0 | 0 | 0 | 0 |
| Reptile | 3 | 3.3 | 0 | 0 | 1 | 1.37 | 0 | 0 | 0 | 0 |
| Unknown | 8 | 8.8 | 2 | 1.4 | 0 | 0 | 0 | 0 | 2 | 0.7 |
| Unknown mammal | 1 | 1.1 | 0 | 0 | 0 | 0 | 0 | 0 | 0 | 0 |
| Totals | 44 | 100 | 109 | 100 | 72 | 100 | 138 | 100 | 283 | 100 |

The precision (one standard deviation of standards) of the δ^15^N and δ^13^C reads was 0.097 and 0.080 respectably, n=34.

### Niche breadth

Pterygoplichtys sp. was the main identifiable prey item in all samples from the Usumacinta basin. Occurrence of ACF in scat samples was highest (49%) in samples collected from Pasion River 7 years after the first report of the catfish, an increase from 9.9% in 2010 (Table 3). ACF occurrence also increased in the San Pedro River, but less than in the Pasion River. O. aureus was an important item (percentage of occurrence > 5%) for otters in the Pasion and San Pedro rivers in 2015 and the Mopan River in 2016 (Table 3).

Based on species accumulation curves, the expected number of prey species was marginally lower in 2015 than in 2010 in Pasion River samples (Fig 2A); no difference was seen for San Pedro River samples (Fig 2B). When all three rivers were compared based on data from 2015-2016, otters from the San Pedro River were expected to have more prey species, those from the Pasion River fewer species, and those from the Mopan River were expected to have a middle number of prey species. Confidence intervals around expected numbers were wide and overlapped, especially between curves from the Mopan River and the other two rivers (Fig 2C). Further, the assumption that all samples used to construct the accumulation curves were independent may have been violated because some of the scats were collected from the same latrine.

**Fig 2.**
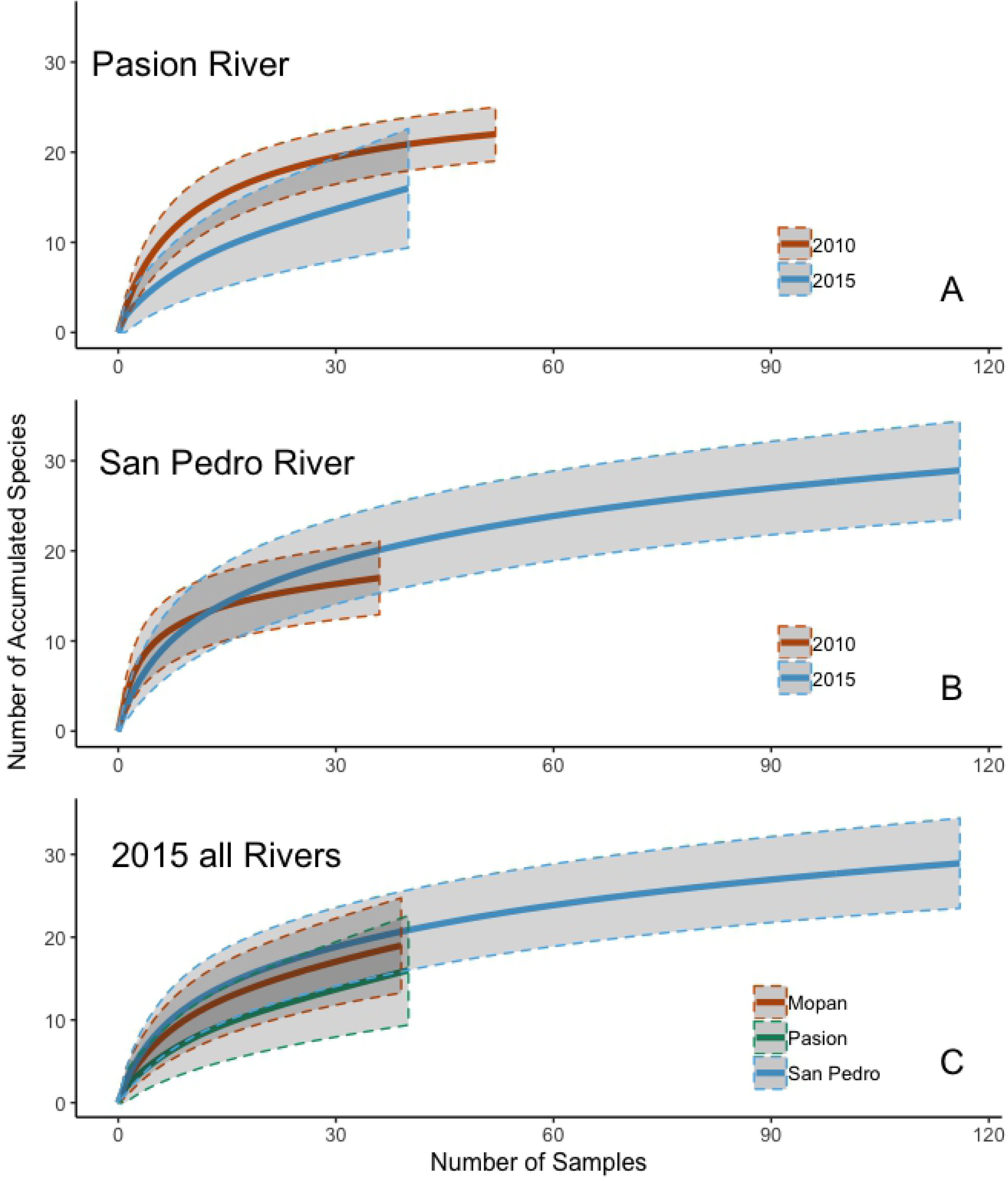
Species accumulation curves for prey species found in scats of Neotropical river otter in the **(A)** Pasion River, Guatemala 2010 (Pa10) and 2015 (Pa15); **(B)** San Pedro River, Guatemala, in 2009 (Sp09) and 2015 (Sp15); and **(C)** Mopan River 2016 (Mo16), Pasion River 2015 (Pa15) and San Pedro River 2015 (Sp15).

Niche breadth (Levin’s index, B_a_) of the Neotropical river otter was lower 7 years after the ACF invasion when compared to 2 years after the invasion in the San Pedro River (B_a_ = 0.53 in 2009 vs 0.29 in 2015). A similar situation was found in Pasion River (B_a_ = 0.47 in 2010 vs 0.18 in 2015). NRO niche breadth varied among the three rivers in 2015, with similar values in San Pedro River and Mopan River and lower values in Pasion River (B_a_ = 0.29, 0.28 and 0.18 respectively; Table 4).

**Table 4.** Neotropical river otter niche breadth (Levin’s index, B_a_) in the study area.

| River | year | Ba | 2.5% quantile | 97.5% quantile |
| --- | --- | --- | --- | --- |
| Mopan | 2016 | 0.29 | 0.36 | 0.23 |
| San Pedro | 2009 | 0.53 | 0.6 | 0.5 |
| San Pedro | 2015 | 0.29 | 0.33 | 0.25 |
| Pasion | 2010 | 0.47 | 0.59 | 0.39 |
| Pasion | 2015 | 0.18 | 0.25 | 0.11 |

Quantiles estimated using 1,000 bootstrap randomizations.

Isotope values ranged from 5.89 to 16.39 δ^15^N and −38.31 to −20.61 δ^13^C (Fig 3) and did not depart from a normal distribution so no transformations were needed. Variance of δ^15^N signatures from fecal samples differed among groups (Levene’s test for homoscedasticity; W = 2.54, *p* = 0.042; Fig 4A). Based on pairwise comparisons, variance of δ^15^N signatures from the Pasion River did not differ significantly between years (σ^2^ = 2.45 in 2010 and σ^2^ = 1.80 in 2015; W = 0.78, *p* = 0.37; Fig 4A). In contrast, variance of δ^15^N differed significantly between years in samples from San Pedro River (σ^2^ = 4.83 in 2009 and σ^2^ = 1.73 in 2015; W = 6.68, *p* < 0.01; Fig 4A). The δ^13^C variances also differed among groups (W = 3.23, *p* < 0.01; Fig 4B), with pairwise contrasts indicating that δ^13^C variances increased across years for Pasion River (σ^2^ = 3.65 in 2010 and σ^2^ = 6.49 in 2015; W = 3.83, *p* = 0.05; Fig 4B) and San Pedro River (σ^2^ = 2.04 in 2009 and σ^2^ = 7.09 in 2015; W = 6.75, *p* = 0.01; Fig 4B).

**Fig 3.**
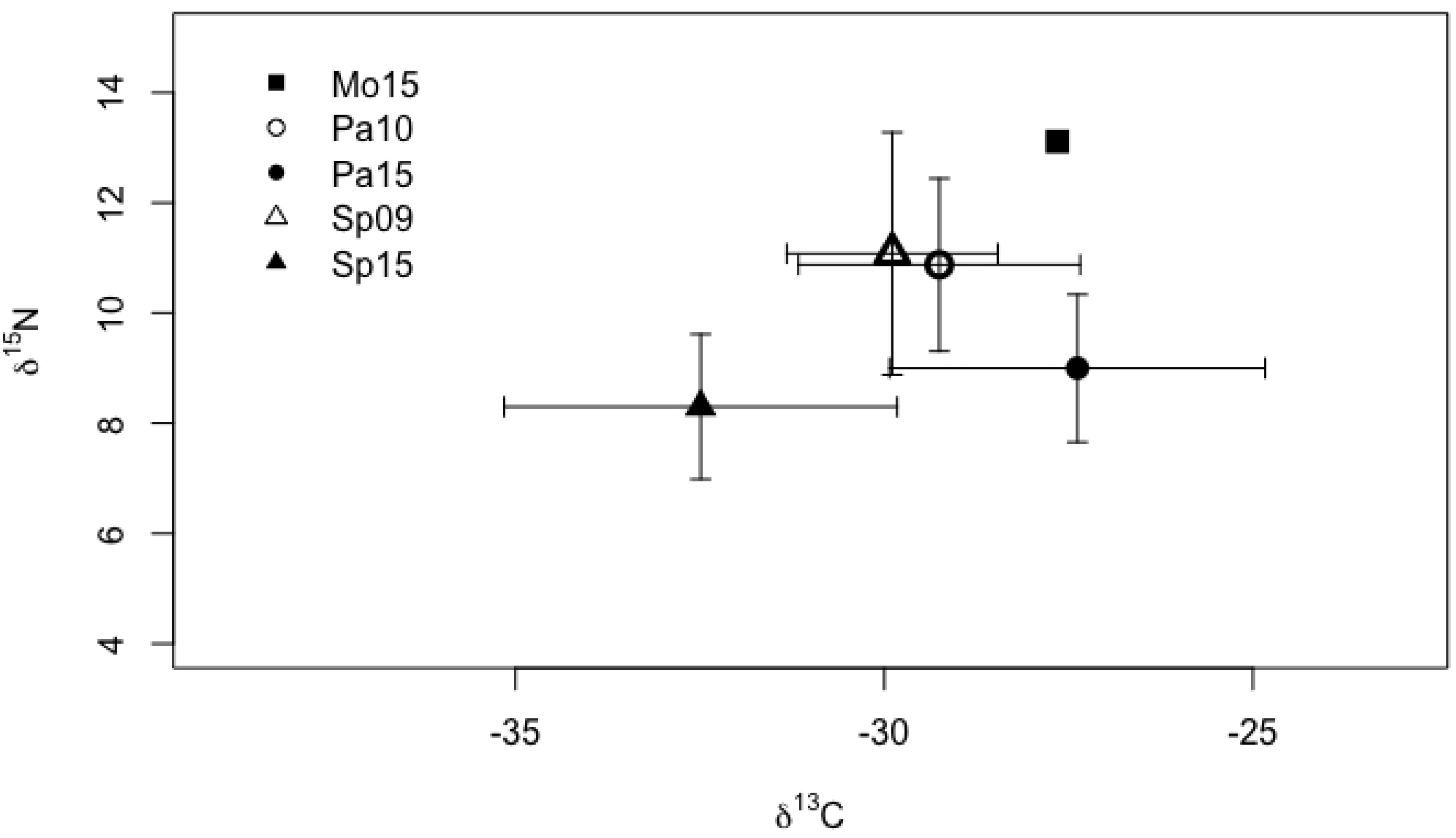
Isotopic values of *δ*^15^N and *δ*^13^C from Neotropical river otter scats collected from the study area. Error bars are one sd. Mo15= samples from Mopan River 2015 (n=1); Pa10, Pa15 = samples from Pasion River 2010 and 2015 (n = 36 in 2010 and 34 in 2015); Sp09, Sp15 = samples from San Pedro River 2009 and 2015 (n = 20 in 2010 and 55 in 2015).

**Fig 4.**
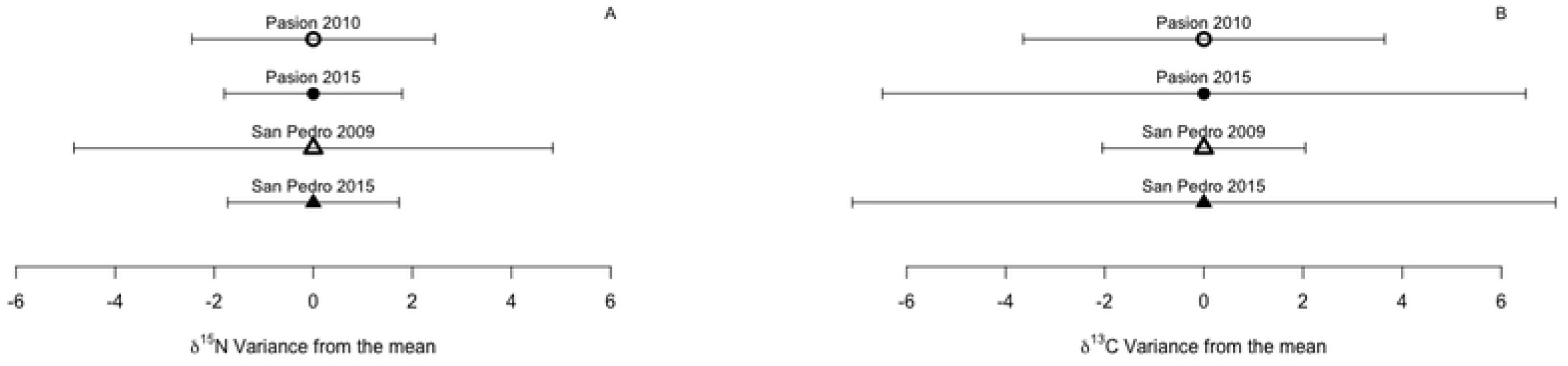
Isotopes Variance from the mean in fecal samples from Neotropical river otters for (A) *δ*^15^N and (B) *δ*^13^C. The mean is set to 0 to help visualize the magnitude of the variances. Pasion River n = 36 in 2010 and 34 in 2015; San Pedro River n = 20 in 2010 and 55 in 2015.

### Trophic level

Calculations of FTL values excluded information from *Maskaheros argenteus* (found in one sample from the Pasion River), insects, reptiles and unknown species because no data on the FTL of those prey items were available. Similarly, crabs and crayfish were excluded because they consume items from different trophic levels and their specific diets are not known for the study area, although crabs were important prey items in Mopan 2016 and Pasion 2010. The highest FTL values for NRO came from the Mopan River in 2016, Pasion River in 2010, and San Pedro River in 2009 with lower values from the Pasion and San Pedro rivers in 2015 (Table 5).

**Table 5.** Neotropical river otter fractional trophic level (FTL) in the study area.

| River | year | FTL | 2.5% quantile | 97.5% quantile |
| --- | --- | --- | --- | --- |
| Mopan | 2016 | 3.98 | 4.11 | 3.83 |
| San Pedro | 2009 | 3.71 | 3.79 | 3.64 |
| San Pedro | 2015 | 3.47 | 3.53 | 3.41 |
| Pasion | 2010 | 3.78 | 3.9 | 3.68 |
| Pasion | 2015 | 3.48 | 3.61 | 3.37 |

Quantiles estimation using 1,000 bootstrap randomizations.

Values of δ^15^N from NRO samples were highest in the Mopan River in 2015 (based on only one specimen), followed by mean values from the Pasion River in 2010 and the San Pedro River in 2009 (Fig 5). Lowest values came from the Pasion and San Pedro rivers in 2015 (Fig 5). Values of δ^15^N from sites in the Usumacinta basin differed across years (ANOVA, F_1,141_ = 67.98; *p* < 0.001) and across rivers (ANOVA, F_1,141_ = 15.53; *p* < 0.001) with no interaction between the two factors (ANOVA, F_1,141_ = 2.76; *p* = 0.10). Higher values were found from scats collected during the early sampling years in the Pasion and San Pedro rivers, two years after the first report of the ACF (post-hoc pairwise *t*-test with Bonferroni adjusted *p*-values: Pasion 2010 vs. 2015 *t* = 5.37, df = 68, *p* < 0.001; San Pedro 2009 vs. 2015, *t* = 5.31, df = 24.122, *p* < 0.001). Mean values of NRO δ^15^N did not differ between the Pasion and San Pedro rivers from same sampling years (Pasion vs. San Pedro 2010-2009 *t* = −0.40, df = 54, *p* = 1.0; Pasion vs. San Pedro 20015, *t* = 2.42, df = 87 *p* = 0.23).

**Fig 5.**
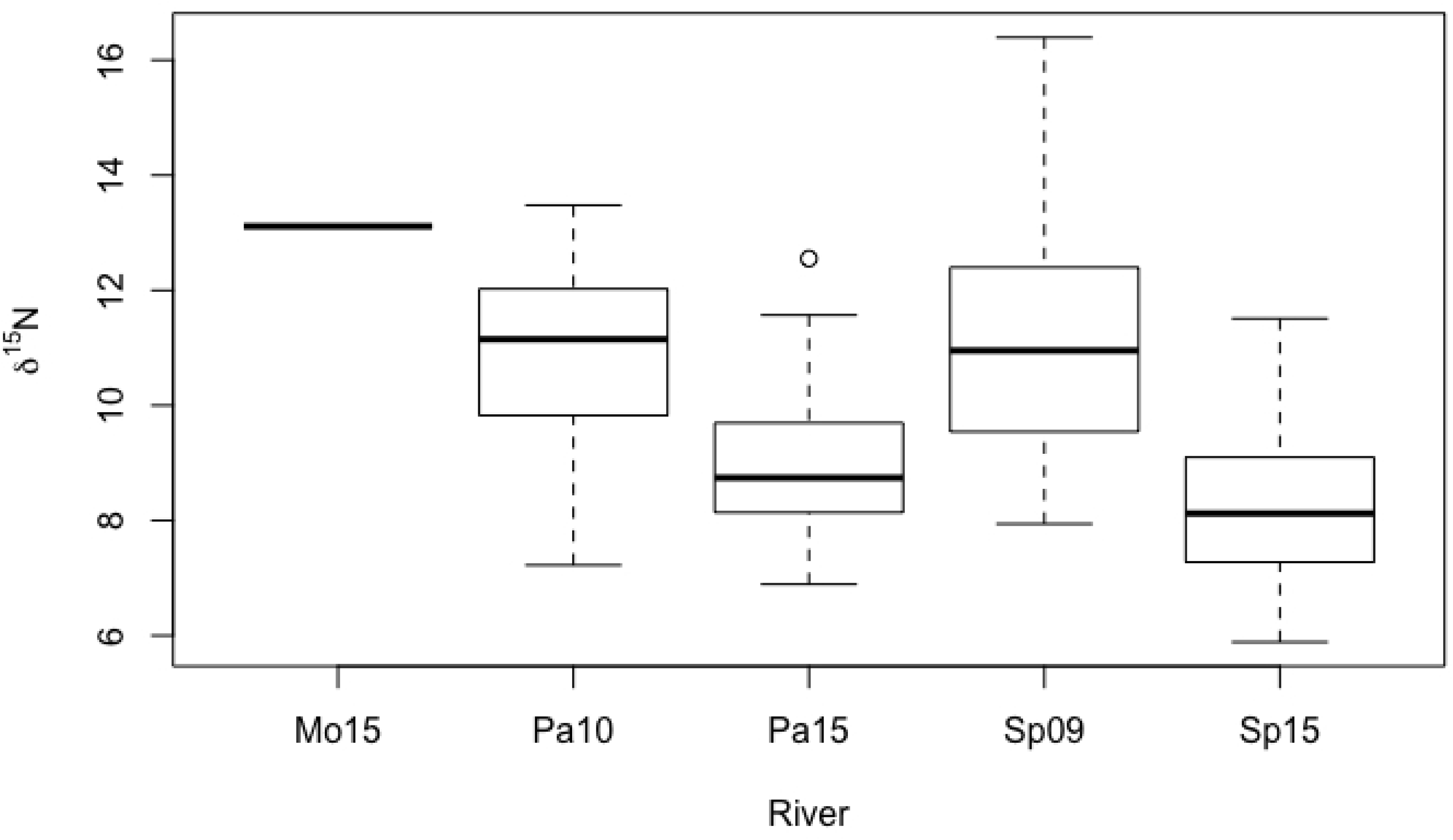
Boxplots for *δ*^15^N in fecal samples from Neotropical river otters in Guatemala. Mo16 = samples from Mopan River 2016 (n = 1); Pa10, Pa15 = samples from Pasion River 2010 and 2015 (n = 36 in 2010 and n = 34 in 2015); Sp09, Sp15 = samples from San Pedro River 2009 and 2015 (n = 20 in 2009 and n = 55 in 2015).

## Discussion

Concordance between the gross scat analysis and stable isotope analysis values strongly supports the idea that an increase in consumption of the armored catfish reduced the dietary niche breadth of the neotropical river otter and trophic level at which the otter feeds in northern Guatemala. As predicted, ACF became the main prey species for the NRO in invaded rivers and, consequently, NRO δ^15^N variances and mean values decreased over time in both invaded rivers (with a weaker decline in Pasion River). The same pattern was observed in the standardized niche breadth index (B_a_). Further, the wider niche breadth (B_a_ values) in the San Pedro River may be related to its higher environmental integrity (located adjacent to a national park) that could help sustain the richness of native NRO prey or reduce the invasiveness of the ACF. This conclusion is supported by the species accumulation curves. Invasive species are predicted to have better chances of establishment in native assemblages that are depleted or disrupted and more likely to have long-term success in systems highly altered by human activity [81]. The increase in δ^13^C variation over time suggests that the NRO diet has included a prey that consumes different producer types or a prey that consumes producers in a different proportion, likely because of the ability of ACF to exploit a different range/proportion of plant resources than natives from the same trophic guild [82]. Furthermore, the decrease in FTL across rivers (Mopan River showing similar values to San Pedro River and higher than Pasion River) combined with lower mean values of δ^15^N provide evidence of a reduction in the NRO trophic level associated with ACF presence.

The range of prey types exploited by NRO changed after the invasion of ACF, with the lowest dietary niche breadth found in Pasion River seven years after the invasion. The dietary NRO niche breadth changed from that of a mild generalist to that of a specialist (0.6 > B_a_ > 0.4 to B_a_ < 0.4, Levin’s standardized index) in Pasion and San Pedro rivers, even though the number of prey species consumed by NRO was highest in the San Pedro River in 2015. This result is concordant with the idea that a specialist can use a wide range of resources but still concentrate on a subset of those resources [83]. It also supports the statement that NRO prey mostly on slow-moving and territorial prey species [36]; the main prey species for NRO in this study included Loricariidae, Cichlidae, large Poeciliidae, and crabs (Table 3).

Results based on GSA and δ^15^N variances were similar for both indexes, with narrower niche breadth 7 years after initiation of the ACF invasion compared to 2 years after the invasion. The narrower dietary niche breath found in the Pasion River in all situations and with both indexes in relation to the San Pedro River supports the idea that the Pasion River prey community was already depleted before the arrival of the ACF, and that the Laguna del Tigre National Park provided some type of protection to the San Pedro River. A similar result was seen in a Bahamas mangrove system for grey snapper (*Lutjanus griseus*) with a reduced niche breadth (based on SIA) found in disturbed areas [5]. Therefore, it is possible that the higher values of NRO niche breadth in San Pedro River in relation to Pasion River are related to differences in the resilience of the two rivers due to differences in habitat conservation. Disturbances may facilitate the ACF or depress populations of native fish. For example, in the Guadalquivir marshes of southwestern Spain, the Eurasian otter (*Lutra lutra*) included high levels of an invasive species (75%; North American red swamp crayfish, *Procambarus clarkii*) in its diet within 10 years of the invasion. In the same area, various waterbirds similarly consumed this invasive species at higher rates in disturbed locations than in natural marshes [1].

In contrast to results from δ^15^N, variances of δ^13^C in fecal samples were greater seven years after the ACF invasion compared to two years after the first sighting. Values of δ^13^C represent the plant source of a food chain and a wider variance may indicate that primary consumers exploit a greater range of producers. Loricariidae may exploit a diverse variety of basal sources or a portion that the natives does not exploit, which may help explain the increase in the variance of δ^13^C in NRO scats, given the increased presence of ACF in the NRO diet [84].

Native predators may act to reduce invasive species numbers [85,86], and such predation could be one of the main biological drivers by which streams resist the invasion of exotic species [87]. Further, predators from different taxa often adapt to and benefit from the consumption of invasive species [3,4,88]. In this context, NRO may act as a buffer to hold ACF populations at low levels and minimize their potential negative effects on the system. The question that arises from this situation, as in other systems where an invasive species becomes the main prey of a native predator [1], is whether the consumption of ACF by NRO and other native predators can facilitate the predators [89]. Greater predator populations might increase depredation on native prey that are threatened by overexploitation or habitat loss [90]. This effect is a valid concern in our study area, where cichlids, a group of fish that is highly appreciated by the local artisanal fisheries [91] were exploited as a group without much change when the consumption of ACF increased (Table 3). Also, concern for the increase of negative interactions between native predator and humans becomes relevant when wild predators establish dense populations in or near human-dominated areas [92,93].

Both GSA and δ^15^N values indicated a reduction in the trophic level at which otters feed in rivers where ACF are present in northern Guatemala. Based on GSA, there were reductions in the FTL of NRO of approximately 0.33 FTL in the Pasion River and 0.2 FTL in the San Pedro River. These reductions may not represent much ecological difference. GSA may, however, under-estimate the consumption of some species and over-estimate the consumption of others either because of differences in digestibility of prey or because we measured presence of prey remains rather than consumed biomass, regardless of the amount of remains (not all remains were identifiable; e.g., spines). In contrast to GSA, SIA may give a more accurate result. Differences in mean δ^15^N were as great as 1.88‰ for Pasion River and 2.78‰ for San Pedro River. If we use the widely accepted 3.4‰ enrichment (Δ^15^N) per FTL, these differences in mean δ^15^N may represent changes of 0.5 to 0.8 FTLs in the Pasion and San Pedro rivers, respectively. The 3.4‰ Δ^15^N value has, however, been criticized. Models and empirical data have shown that this enrichment factor can underestimate FTL of marine predators [94]. In any case, the observed mean δ^15^N values for NRO in both the Pasion and San Pedro rivers apparently represents a decrease in trophic level.

A reduction in the trophic level at which otters feed can have diverse effects on the riverine ecosystem. These effects may be difficult to anticipate and can compete with or interact with each other. It could mean predator release for other prey species that would benefit from reduced predation pressure [95,96]. On the other hand, consumption of the invasive species may benefit the predator, eventually leading to higher predator densities that could increase pressure on other native species. A model evaluating this situation suggests that predation on native prey by a native predator whose numbers have been enhanced by consumption of an invasive species can be more harmful than direct competition between native and invasive species [97]. Empirical data using SIA for golden eagles (Aquila chrysaetos) suggests that these eagles colonized the California Channel Islands after the introduction of feral pigs (Sus scrofa) [90]. Nonetheless, eagles still preyed on endemic meso-carnivores, including a fox (Urocyon littoralis) and skunk (Spilogale gracilis amphiala), pushing the fox towards extinction [90].

Another potential effect that needs to be evaluated is the reduction of trophic levels in the system by moving energy directly from primary consumers to top predators. This results can occur by eliminating food-web links in the mid-trophic levels through competition or predation facilitated by a numerical response of predators in response to the high abundance of the invasive [1]. A similar situation was found in the United Kingdom, where researchers compared the fish assemblage in a pond with a low-trophic-level invasive cyprinid (*Pseudorasbora parva*) composing > 99% of fish present to that in another pond without the cyprinid. They reported a reduction in the δ^15^N values of piscivorous fish and a mean reduction in the δ^15^N of the complete fish community [98]. Further studies are needed to investigate the effect of different types of land management, as well as factors that indicate the ecological integrity of communities, on the ability of communities to resist or facilitate the invasion of exotic species and their interactions with native predators.

## Acknowledgments

We wish to acknowledge the institutions and all the people who helped us through those institutions to make the fieldwork and laboratory work possible: Eli.S.A., WCS-Guatemala, BALAM, CONAP-Peten, Propeten, Defensores de la Naturaleza Peten, FUNDAECO, CECON, the Univeridad de San Carlos de Guatemala Escuela de Biología, the University of Florida Departement of Geological Sciences Light Stable Isotopes Mass Spectrometry laboratory, Roan Balas, Byron Castellanos, Yobany Tut, Julio Madrid, Rony Garcia, Rafael Cevallos, Mercedes Barrios, Rosalito Barrios, Manuel Lepe, Werner Paz, Silja Ramirez, Leonel Zisse, Marta Pujol and Jason Curtis. The people who helped in the data gathering: Alejandro Chen, Alejandro Mérida, Alfredo Choc, Belarmino García, Carlos Cifuentes, Dastin Ramírez, Elder Godoy, Elmer Monzón, Ervin Flores, Fredy Tot, Jeovany Nolberto Tut Pacheco, René de Jesús Mauricio Te, Ricardo Coc Caal, Samuel Yatz, Juan Rodas but specially to Andrea Paiz and Francisco Cordova who were present during most of the sampling occasions in 2015-2016 and Yasmín Quintana in 2010. Also, to the people whose comments, reviews, and advice helped to improve this document: Christina Romagosa, Bill Pine, Harry Jones, Laura Gelin, Jose Soto, Claudio Moraga, Audrey Wilson, Yasmín Quintana, Flavia Montalvo, Farah Carrasco, and Jason Curtis.

## References

1. Tablado Z, Tella JL, Sánchez-Zapata JA, Hiraldo F. The paradox of the long-term positive effects of a North American crayfish on a European community of predators. Conserv Biol. 2010;24: 1230–1238. doi:10.1111/j.1523-1739.2010.01483.x

2. Cattau CE, Fletcher Jr RJ, Kimball RT, Miller CW, Kitchens WM. Rapid morphological change of a top predator with the invasion of a novel prey. Nat Ecol Evol. Springer US; 2017; 1–8. doi:10.1038/s41559-017-0378-1

3. Cattau CE, Fletcher Jr. RJ, Reichert BE, Kitchens WM. Counteracting effects of a nonnative prey on the demography of a native predator culminate in positive population growth. Ecol Appl. 2016;26: 1952–1968. doi:10.1890/15-1020.1

4. King RB, Ray JM, Stanford KM. Gorging on gobies: Beneficial effects of alien prey on a threatened vertebrate. Can J Zool. 2006;84: 108–115. doi:10.1139/z05-182

5. Layman CA, Quattrochi JP, Peyer CM, Allgeier JE. Niche width collapse in a resilient top predator following ecosystem fragmentation. Ecol Lett. 2007;10: 937–944. doi:10.1111/j.1461-0248.2007.01087.x

6. Fricke R, Eschmeyer WN, Van Der Laan R. Catalog of Fishes. In: 04-September-2018 [Internet]. 2018 [cited 6 Sep 2018]. doi:10.11646/zootaxa.3882.1.1

7. Nico LG, Jelks HL, Tuten T. Non-native suckermouth armored catfishes in Florida: Description of nest burrows and burrow colonies with assessment of shoreline conditions. Aquat Nuis Species Res Bull. 2009;9: 1–30.

8. Capps K a., Nico LG, Mendoza-Carranza M, Arévalo-Frías W, Ropicki AJ, Heilpern S a., et al. Salinity tolerance of non-native suckermouth armoured catfish (Loricariidae: Pterygoplichthys) in southeastern Mexico: Implications for invasion and dispersal. Aquat Conserv Mar Freshw Ecosyst. 2011;21: 528–540. doi:10.1002/aqc.1210

9. da Cruz AL, da Silva HR, Lundstedt LM, Schwantes AR, Moraes G, Klein W, et al. Air-breathing behavior and physiological responses to hypoxia and air exposure in the air-breathing loricariid fish, Pterygoplichthys anisitsi. Fish Physiol Biochem. 2013;39: 243–256. doi:10.1007/s10695-012-9695-0

10. Stevens PW, Blewett DA, Casey JP. Short-term effects of a low dissolved oxygen event on estuarine fish assemblages following the passage of hurricane Charley. Estuaries and Coasts. 2006;29: 997–1003. doi:10.1007/BF02798661

11. Corcuera Zabarburú CA. Evaluación de la temperatura letal máxima para Hypostomus hemicochiodon y Pterygoplichthys pardalis en el centro de investigaciones Carlos Miguel Castañeda Ruiz. Universidad nacional Toribio Rodríguez de Mendoza de Amazonas. 2015.

12. de Oliveira C, Taboga S. R, Smarra AL, Bonilla-Rodriguez GO. Microscopical aspects of accessory air breathing through a modified stomach in the armoured catfish Liposarcus anisitsi (Siluriformes, Loricariidae). Cytobios. 2001;105: 153–162.

13. Froese R, Pauly D. FishBase. In: 06/2018 [Internet]. 2018 [cited 6 Sep 2018]. Available: www.fishbase.org

14. Gibbs MA, Shields JH, Lock DW, Talmadge KM, Farrell TM. Reproduction in an invasive exotic catfish Pterygoplichthys disjunctivus in Volusia Blue Spring, Florida, U.S.A. J Fish Biol. 2008;73: 1562–1572. doi:10.1111/j.1095-8649.2008.02031.x

15. Gibbs M a., Kurth BN, Bridges CD. Age and growth of the loricariid catfish Pterygoplichthys disjunctivus in Volusia Blue Spring, Florida. Aquat Invasions. 2013;8: 207–218. doi:10.3391/ai.2013.8.2.08

16. Marchetti MP, Moyle PB, Levine R. Alien fishes in California watersheds: Characteristics of successful and failed invaders. Ecol Appl. 2004;14: 587–596. doi:10.1890/02-5301

17. Orfinger AB, Goodding DD. The global invasion of the suckermouth armored catfish genus pterygoplichthys (Siluriformes: Loricariidae): Annotated list of species, distributional summary, and assessment of impacts. Zool Stud. 2018;57: 1–16. doi:10.6620/ZS.2018.57-07

18. Wakida-Kusunoki AT, Ruiz-Carus R, Amador-del-Angel E. Amazon sailfin catfish, Pterygoplichthys pardalis (Castelnau, 1855) (Loricariidae), another exotic species established in Southeastern México. Southwest Nat. 2007;52: 141–144. doi:10.1894/0038-4909(2007)52[141:ASCPPC]2.0.CO;2

19. Wakida-Kusunoki AT, Amador-Del-Angel LE. Nuevos registros de los plecos Pterygoplichthys pardalis (Castelnau 1855) y P. disjunctivus (Weber 1991) (Siluriformes: Loricariidae) en el Sureste de México. Hidrobiologica. 2008;18: 251–255.

20. Hubbs C, Edwards RJ, Garrett GP. An annotated checklist of the freshwater fishes of Texas, with keys to identification of species. Second Edi. Austin: Texas Academy of Science; 2008.

21. Wu L-W, Liu C, Lin S-M. Identification of exotic sailfin catfish species (Pterygoplichthys, Loricariidae) in Taiwan based on morphology and mtDNA sequences. Zool Stud. 2011;50: 235–246.

22. Capps KA, Flecker AS. Invasive fishes generate biogeochemical hotspots in a nutrient-limited system. PLoS One. 2013;8: e54093. doi:10.1371/journal.pone.0054093

23. Capps KA, Flecker AS. High impact of low-trophic-position invaders: Nonnative grazers alter the quality and quantity of basal food resources. Freshw Sci. 2015;34: 784–796. doi:10.1086/681527.

24. Bunkley-Williams L, Williams Jr. EH, Lilystrom CG, Corujo-Flores I, Zerbi AJ, Aliaume C, et al. The South American sailfin armored catfish, Liposarcus multiradiatus (Hancock), a new exotic established in Puerto Rican fresh waters. Caribb J Sci. 1994;30: 90–94.

25. Chaichana R, Pouangcharean S, Yoonphand R. Foraging effects of the invasive alien fish Pterygoplichthys on eggs and first-feeding fry of the native Clarias macrocephalus in Thailand. Kasetsart J. 2013;47: 581–588.

26. Hubilla M, Kis F, Primavera J. Janitor fish Pterygoplichthys disjunctivus in the Agusan Marsh: A threat to freshwater biodiversity. J Environ Sci Manag. 2007;10: 10–23.

27. Capps KA, Ulseth A, Flecker AS. Quantifying the top-down and bottom-up effects of a non-native grazer in freshwaters. Biol Invasions. 2014;17: 1253–1266. doi:10.1007/s10530-014-0793-z

28. Nico LG, Loftus WF, Reid JP. Interactions between non-native armored suckermouth catfish (Loricariidae: Pterygoplichthys) and native Florida manatee (Trichechus manatus latirostris) in artesian springs. Aquat Invasions. 2009;4: 511–519. doi:10.3391/ai.2009.4.3.13

29. Nico LG. Nocturnal and diurnal activity of armored suckermouth catfish (Loricariidae: Pterygoplichthys) associated with wintering Florida manatees (Trichechus manatus latirostris). Neotrop Ichthyol. 2010;8: 893–898. doi:10.1590/S1679-62252010005000014

30. Gibbs M, Futral T, Mallinger M, Martin D, Ross M. Disturbance of the Florida manatee by an invasive catfish. Southeast Nat. 2010;9: 635–648. doi:10.1656/058.009.0401

31. Lienart G-DH, Rodiles-Hernández R, Capps KA. Nesting Burrows and Behavior of Nonnative Catfishes (Siluriformes: Loricariidae) in the Usumacinta-Grijalva Watershed, Mexico. Southwest Nat. 2013;58: 238–243. doi:10.1894/0038-4909-58.2.238

32. Toro-Ramírez A, Wakida-Kusunoki AT, Amador-del Ángel LE, Cruz-Sánchez JL. Common snook [Centropomus undecimalis (Bloch, 1792)] preys on the invasive Amazon sailfin catfish [Pterygoplichthys pardalis (Castelnau, 1855)] in the Palizada River, Campeche, southeastern Mexico. J Appl Ichthyol. 2014;30: 532–534. doi:10.1111/jai.12391

33. Ríos-Muñoz CA. Depredación de pez diablo (Loricariidae: Pterygoplichthys) por el cormorán oliváceo (Phalacrocorax brasilianus) en Villahermosa, Tabasco, México. Huitzil. 2015;16: 62–65.

34. Kruuk H, Balharry E, Taylor PT. Oxygen consumption of the Eurasian Otter Lutra lutra in relation to water temperature. Physiol Zool. 1994;67: 1174–1185.

35. Pfeiffer P, Culik BM. Energy metabolism of underwater swimming in river-otters (Lutra lutra L.). J Comp Physiol – B Biochem Syst Environ Physiol. 1998;168: 143–148. doi:10.1007/s003600050130

36. Rheingantz ML, Oliveira-santos LG, Waldemarin HF, Caramaschi EP. Are otters generalists or do they prefer larger, slower prey? Feeding flexibility of the neotropical otter Lontra longicaudis in the Atlantic forest. IUCN Otter Spec Gr Bull. 2012;29: 80–94.

37. Chemes SB, Giraudo AR, Gil G. Dieta de Lontra Longicaudis (Carnivora, Mustelidae) en el Parque Nacional El Rey (Salta, Argentina) y su comparación con otras poblaciones de la cuenca. Mastozoología Neotrop. 2010;17: 19–29. doi:10.2307/3504393

38. Kasper CB, Bastazini VAG, Salvi J, Grillo HCZ. Trophic ecology and the use of shelters and latrines by the Neotropical otter (Lontra longicaudis) in the Taquari Valley, Southern Brazil. Iheringia Série Zool. 2008;98: 469–474.

39. Mayor-Victoria R, Botero-Botero Á. Dieta de la nutria neotropical Lontra longicaudis (Carnívora, mustelidae) en el Río Roble, Alto Cauca, Colombia. Acta Biológica Colomb. 2010;15: 237–244.

40. Silva FA Da, Nascimento EDM, Quintela FM. Diet of Lontra longicaudis (Carnivora: Mustelidae) in a pool system in Atlantic forest of Minas Gerais State, southeastern Brazil. Acta Sci Biol Sci. 2012;34: 407–412. doi:10.4025/actascibiolsci.v34i4.10332

41. Casariego-Madorell A, List R, Ceballos G. ASPECTOS BÁSICOS SOBRE LA ECOLOGÍA DE LA NUTRIA DE RÍO (Lontra longicaudis annectens) PARA LA COSTA DE OAXACA. Rev Mex Mastozoología. 2006;10: 71–74.

42. Monroy-Vilchis O, Mundo V. Nicho trófico de la nutria neotropical (Lontra longicaudis) en un ambiente modificado, Temascaltepec, México. Rev Mex Biodivers. 2009;80: 801–806.

43. Rheingantz ML, Waldemarin HF, Rodrigues L, Moulton TP. Seasonal and spatial differences in feeding habits of the Neotropical otter Lontra longicaudis (Carnivora: Mustelidae) in a coastal catchment of southeastern Brazil. Zoologia. 2011;28: 37–44. doi:10.1590/S1984-46702011000100006

44. Marques Quintela F, Almeida Porciuncula R, Pinto Colares E. Dieta de Lontra longicaudis (Olfer) (Carnivora, Mustelidae) em um arroio costeiro da região sul do Estado do Rio Grande do Sul, Brasil. Neotrop Biol Conserv. 2008;3: 119–125. doi:10.4013/nbc.20083.03

45. Sousa KS, Saraiva DD, Colares EP. Intra-annual dietary variation in the neotropical otter from southern Brazil. Mammal Study. 2013;38: 155–162.

46. Kasper CB, Feldens MJ, Salvi J, César H, Grillo Z. Estudo preliminar sobre a ecologia de Lontra longicaudis (Olfers) (Carnivora, Mustelidae) no Vale do Taquari, Sul do Brasil. Rev Bras Zool. 2004;21: 65–72.

47. Pardini R. Feeding ecology of the neotropical river otter Lontra longicaudis in an Atlantic Forest stream, south-eastern Brazil. J Zool. 1998;245: 385–391. doi:10.1111/j.1469-7998.1998.tb00113.x

48. Holdridge LR, Grenke WC. Forest environments in tropical life zones: a pilot study. 1971.

49. Willink PW, Barrientos C, Kihn HA, Chernoff B. An ichthyological survey of Laguna del Tigre National Park, Peten, Guatemala. In: Bestelmeyer BT, Alonso LE, editors. A biological assessment of Laguna del Tigre National Park, Petén, Guatemala RAP bulleting of biological assessment 16. Washintong, DC: Conservation International; 2000. pp. 41–48.

50. Granados-Dieseldorff P, Christensen MF, Kihn-Pineda PH. Fishes from Lachuá lake, upper Usumacinta basin, guatemala. Check List. 2012;8: 95–101.

51. Greenfield DW, Thomerson JE. Fishes of the continental waters of Belize. Fishes of the continental waters of Belize. University Press of Florida; 1997.

52. Aranda-Sánchez JM. Manual para el rastreo de mamíferos silvestres de México. Primera ed. Suárez Huesca HK, Hernández Vázquez L, Escobar Ramírez F de J, editors. México D.F.: Ciba Diseño y Arte Editorial; 2012.

53. Kasper CB, Salvi J, Zanardi Grillo HC. Estimativa do tamanho de duas espécies de ciclídeos (Osteichthyes, Perciformes) predados por Lontra longicaudis (Olfers) (Carnivora, Mustelidae), através de análise das escamas. Rev Bras Zool. 2004;21: 499–503.

54. Juárez-Sánchez D. Scale guide to identify medium and large freshwater fish from northern Guatemala [Internet]. University of Florida. 2017. Available: http://ufdc.ufl.edu/l/IR00009693/00001

55. Juárez-Sánchez D, Estrada C, Bustamante M, Moreira J, Quintana Y, López J. Guia Ilustrada de pelos para la identificación de mamiferos mayores y medianos de Guatemala. 2 da edici. Dirección General de Investigación (DIGI), editor. Guatemala; 2010.

56. Colwell RK, Chang XM, Chang J. Interpolating, extrapolating, and comparing incidence-based species accumulation curves. Ecology. 2004;85: 2717–2727. doi:10.1890/03-0557

57. Levins R. Evolution in changing enviroment: Some theoretical explorations. Prinston university press. Princeton, New Jersey; 1968.

58. Krebs CJ. Ecological Methodology. Second Edi. New York: Addison-Wesley Educational Publishers, Inc.; 1999. doi:10.1037/023990

59. Novakowski GC, Hahn NS, Fugi R. Diet seasonality and food overlap of the fish assemblage in a pantanal pond. Neotrop Ichthyol. 2008;6: 567–576. doi:10.1590/S1679-62252008000400004

60. Pauly D, Palomares M. Fishing down marine food web: It is far more pervasive than we thought. Bull Mar Sci. 2005;76: 197–211.

61. Post DM. Using stable isotopes to estimate trophic position: Models, methods, and assumptions. Ecology. 2002;83: 703–718. doi:Doi 10.2307/3071875

62. Vander Zanden MJ, Casselman JM, Rasmussen JB. Stable isotope evidence for the food web consequences of species invasions in lakes. Nature. 1999;401: 464–467.

63. Petersen BJ, Fry B. Stable Isotopes in Ecosystem Studies. Annu Rev Ecol Syst. 1987;18: 293–320. doi:10.1146/annurev.es.18.110187.001453

64. Fry B. Stable isotope ecology. 3rd editio. United States of America.: Springer Science+Business Media, LLC; 2008. doi:10.1007/0-387-33745-8

65. Kelly JF. Stable isotopes of carbon and nitrogen in the study of avian and mammalian trophic ecology. Can J Zool. 2000;78: 1–27. doi:10.1139/z99-165

66. Blundell G, Ben-David M, Bowyer R. Sociality in river otters: Cooperative foraging or reproductive strategies? Behav Ecol. 2002;13: 134–141. doi:10.1093/beheco/13.1.134

67. Angerbjorn A, Hersteinsson P, Liden K, Nelson E. Dietary variation in arctic foxes (Alopex lagopus): An analysis of stable carbon isotopes. Oecologia. 1994;99: 226–232. doi:10.1007/BF00627734

68. Darimont CT, Reimchen TE. Intra-hair stable isotope analysis implies seasonal shift to salmon in gray wolf diet. Can J Zool. 2002;80: 1638–1642. doi:10.1139/z02-149

69. Aurioles-Gamboa D, Newsome SD, Salazar-Pico S, Koch PL. Stable isotope differences between sea lions (Zalophus) from the Gulf of California and Galapagos Islands. J Mammal. 2009;90: 1410–1420. doi:10.1644/08-mamm-a-209r2.1

70. Fortin JK, Schwartz CC, Gunther KA, Teisberg JE, Haroldson MA, Evans MA, et al. Dietary adjustability of grizzly bears and American black bears in Yellowstone National Park. J Wildl Manage. 2013;77: 270–281. doi:10.1002/jwmg.483

71. Wengeler WR, Kelt D a., Johnson ML. Ecological consequences of invasive lake trout on river otters in Yellowstone National Park. Biol Conserv. 2010;143: 1144–1153. doi:10.1016/j.biocon.2010.02.012

72. Schoeninger MJ, DeNiro MJ, Tauber H. Stable nitrogen isotope ratios of bone collagen reflect marine and terrestrial components of prehistoric human diet. Science (80-). 1983;220: 1381–1383.

73. Newsome SD, Phillips DL, Culleton BJ, Guilderson TP, Koch PL. Dietary reconstruction of an early to middle Holocene human population from the central California coast: Insights from advanced stable isotope mixing models. J Archaeol Sci. 2004;31: 1101–1115. doi:10.1016/j.jas.2004.02.001

74. Salvarina I, Yohannes E, Siemers BM, Koselj K. Advantages of using fecal samples for stable isotope analysis in bats: Evidence from a triple isotopic experiment. Rapid Commun Mass Spectrom. 2013;27: 1945–1953. doi:10.1002/rcm.6649

75. Crowley S, Johnson CJ, Hodder DP. Spatio-temporal variation in river otter (Lontra canadensis) diet and latrine site activity. Écoscience. 2013;20: 28–39. doi:10.2980/20-1-3509

76. Crait JR, Ben-David M. Effects of river otter activity on terrestrial plants in trophically altered Yellowstone Lake. Ecology. 2007;88: 1040–1052. doi:10.1890/06-0078

77. Codron D, Codron J, Lee-Thorp JA, Sponheimer M, de Ruiter D. Animal diets in the Waterberg based on stable isotopic composition of faeces. South African J Wildl Res. 2005;35: 43–52.

78. Hatch KA, Roeder BL, Buckman RS, Gale BH, Bunnell ST, Eggett DL, et al. Isotopic and gross fecal analysis of American black bear scats. Ursus. 2011;22: 133–140. doi:10.2192/URSUS-D-10-00034.1

79. Bearhop S, Adams CE, Waldron S, Fuller RA, Macleod H. Determining trophic niche width: A novel approach using stable isotope analysis. J Anim Ecol. 2004;73: 1007–1012. doi:10.1111/j.0021-8790.2004.00861.x

80. R Core Team. R: A Language and Environment for Statistical Computing [Internet]. Vienna, Austria: R Foundation for Statistical Computing; 2018. Available: https://www.r-project.org/

81. Moyle PB, Light T. Biological invasions of fresh water: Empirical rules and assembly theory. Biol Conserv. 1996;78: 149–161. doi:10.1016/0006-3207(96)00024-9

82. Lopes CA, Manetta GI, Figueiredo BRS, Martinelli LA, Benedito E. Carbon from littoral producers is the major source of energy for bottom-feeding fish in a tropical floodplain. Environ Biol Fishes. 2015;98: 1081–1088. doi:10.1007/s10641-014-0343-7

83. Peers MJL, Thornton DH, Murray DL. Reconsidering the specialist-generalist paradigm in niche breadth dynamics: Resource gradient selection by Canada lynx and bobcat. PLoS One. 2012;7: e51488. doi:10.1371/journal.pone.0051488

84. Burress ED, Duarte A, Gangloff MM, Siefferman L. Isotopic trophic guild structure of a diverse subtropical South American fish community. Ecol Freshw Fish. 2013;22: 66–72. doi:10.1111/eff.12002

85. Ward-Fear G, Brown GP, Shine R. Using a native predator (the meat ant, Iridomyrmex reburrus) to reduce the abundance of an invasive species (the cane toad, Bufo marinus) in tropical Australia. J Appl Ecol. 2010;47: 273–280. doi:10.1111/j.1365-2664.2010.01773.x

86. Wanger TC, Wielgoss AC, Motzke I, Clough Y, Brook BW, Sodhi NS, et al. Endemic predators, invasive prey and native diversity. Proc R Soc B Biol Sci. 2011;278: 690–694. doi:10.1098/rspb.2010.1512

87. Baltz DM, Moyle PB. Invasion resistance to introduced species by a native assemblage of California stream fishes. Ecol Appl. 1993;3: 246–255.

88. Skewes O, Moraga CA, Arriagada P, Rau JR. El jabalí europeo (Sus scrofa): Un invasor biológico como presa reciente del puma (Puma concolor) en el sur de Chile. Rev Chil Hist Nat. 2012;85: 227–232. doi:10.4067/S0716-078X2012000200009

89. Cameron EK, Bayne EM. Invasion by a non-native ecosystem engineer alters distribution of a native predator. Divers Distrib. 2012;18: 1190–1198. doi:10.1111/j.1472-4642.2012.00912.x

90. Roemer GW, Donlan CJ, Courchamp F. Golden eagles, feral pigs, and insular carnivores: How exotic species turn native predators into prey. Proc Natl Acad Sci U S A. 2002;99: 791–796. doi:10.1073/pnas.012422499

91. Barrientos C, Quintana Y, Elías DJ, Rodiles-Hernández R. Peces nativos y pesca artesanal en la cuenca Usumacinta, Guatemala. Mex Rev Mex Biodivers. 2018;89: S118–S130. doi:10.22201/ib.20078706e.2018.4.2180

92. Yirga G, Leirs H, De Iongh HH, Asmelash T, Gebrehiwot K, Deckers J, et al. Spotted hyena (Crocuta crocuta) concentrate around urban waste dumps across Tigray, northern Ethiopia. Wildl Res. 2015;42: 563–569. doi:10.1071/WR14228

93. Bateman PW, Fleming PA. Big city life: Carnivores in urban environments. J Zool. 2012;287: 1–23. doi:10.1111/j.1469-7998.2011.00887.x

94. Hussey NE, Macneil MA, Mcmeans BC, Olin JA, Dudley SFJ, Cliff G, et al. Rescaling the trophic structure of marine food webs. Ecol Lett. 2014;17: 239–250. doi:10.1111/ele.12226

95. Rodriguez LF. Can invasive species facilitate native species? Evidence of how, when, and why these impacts occur. Biol Invasions. 2006;8: 927–939. doi:10.1007/s10530-005-5103-3

96. Grosholz ED, Ruiz GM, Dean C a, Shirley K a, John L, Connors PG, et al. The impacts of a nonindigenous marine predator in a California Bay. Ecology. 2000;81: 1206–1224.

97. Noonburg EG, Byers JE. More harm than good: When invader vulnerabilty to predators enhances impact on native species. Ecology. 2005;86: 2555–2560. doi:10.1890/05-0143

98. Britton JR, Davies GD, Harrod C. Trophic interactions and consequent impacts of the invasive fish Pseudorasbora parva in a native aquatic foodweb: A field investigation in the UK. Biol Invasions. 2010;12: 1533–1542. doi:10.1007/s10530-009-9566-5

